# Physiological and ecological consequences of the water optical properties degradation on reef corals

**DOI:** 10.1101/2021.02.18.431834

**Authors:** Tomás López-Londoño, Claudia T. Galindo-Martínez, Kelly Gómez-Campo, Luis A. González-Guerrero, Sofia Roitman, F. Joseph Pollock, Valeria Pizarro, Mateo López-Victoria, Mónica Medina, Roberto Iglesias-Prieto

## Abstract

Degradation of water optical properties due to anthropogenic disturbances is a common phenomenon in coastal waters globally. Although this condition is associated with multiple drivers that affect corals health in multiple ways, its effect on light availability and photosynthetic energy acquisition has been largely neglected. Here, we describe how declining the water optical quality in a coastal reef exposed to a turbid plume of water originating from a man-made channel compromise the functionality of the keystone coral species *Orbicella faveolata*. We found highly variable water optical conditions with significant effects on the light quantity and quality available for corals. Reduction of light penetration into the water column elicits the development of low-light phenotypes close to theoretical limits of photoacclimation despite their occurrence at shallow depths. Predicted photosynthetic energy depletion with increasing depth is associated with patterns of colony mortality and contraction of the habitable space for the population. A numerical model illustrates the potential effect the progressive degradation of water optical properties on the gradual mortality and population decline of *O. faveolata*. Our findings suggest that preserving the water optical properties seeking to maximize light penetration into the water column may have an extraordinary impact on coral reefs conservation, mostly toward the deeper portions of reefs.

## Introduction

Coral reefs are recognized not only as one of the most complex, productive and biologically diverse ecosystems (Odum and Odum 1955), but also as icons of the devastating effects of anthropogenic stressors to the natural environment. At a planetary scale, we have already lost one fifth of total coral reefs (Wilkinson 2008) as a result of the synergistic effects of large-scale stressors (Ainsworth et al. 2016) and regional and local factors (Carlson et al. 2019). Reefs in the Indo-Pacific and the Caribbean, in particular, have lost nearly half of their coral cover due to human impacts (Bruno and Selig 2007; Jackson 2014). Approximately 60% of remaining coral reef areas are threatened by the continuous increase of sediments, nutrients, and pollutants in the water associated with coastal development and terrestrial runoff (Burke et al. 2011; Carlson et al. 2019).

Ample evidence demonstrates that water-quality deterioration affects reef corals in different ways. Two main stress-related factors are: a) interference of nutrient enrichment in the host’s capacity to control populations of algal symbionts and opportunistic microorganisms (*e*.*g*., Symbiodiniaceae and bacteria) (Shantz and Burkepile 2014); and b) disease prevalence and energetic losses resulting from the physical disturbance of particle abrasion and deposition on coral tissue (Junjie et al. 2014; Pollock et al. 2014). The effects of altered water optical properties and light climate derived from pollution on the physiology and ecology of reef corals has been largely neglected (Yentsch et al. 2002), although this condition has drawn increasing attention (Anthony et al. 2004; Bessell-Browne et al. 2017; Omachi et al. 2019). Current evidence suggests that the energetic imbalance derived from reduced light penetration in the water column compromises the reef-building capacity and coral survivorship, mainly in areas increasingly exposed to terrestrial runoff and dredging-related activities (Junjie et al. 2014; Bessell-Browne et al. 2017; Fisher et al. 2019).

Varadero reef in the southern end of the Caribbean, at the mouth of the Cartagena Bay, Colombia, is exposed to a turbid plume of water originating from a man-made channel (Pizarro et al. 2017). The Dique channel currently delivers large freshwater discharges (currently 8,833 m^-3^ s^-1^) with high sediment load (23,906 t d^-1^) into the Cartagena Bay (Restrepo et al. 2018) as a result of the major rectification works that took place during the 1920’s - 1980’s period (Mogollón 2013). The permanence of this reef under suboptimal environmental conditions provides a unique opportunity to explore the effects of the progressive degradation of the water column optical properties on reef corals. Here, we investigated the effects of these disturbances on the physiology and ecology of the keystone coral species *Orbicella faveolata*, a major reef-builder in the Caribbean. A better understanding of these effects is essential for predicting future impacts associated with coastal development and terrestrial runoff and for the implementation of more effective management efforts seeking to control local key stressors on coral reef ecosystems.

## Material and Methods

### Site selection criteria

Two sites within the Cartagena Reef System with contrasting water optical properties were considered: a turbid-water reef “Varadero”, close to the Dique channel outlet (10° 18’ 23.3’’ N, 75° 35’ 08.0’’ W), and a clear-water reef located 21 km southwest of Varadero within the marine protected area Parque Nacional Natural Corales del Rosario y San Bernardo, hereafter referred as “Rosario” (10° 11’ 12.1’’ N, 75° 44’ 43.0’’ W) (Fig. 1). The Dique channel is a man-made distributary from the Magdalena River, the largest river system of Colombia and major contributor of continental fluxes into the Caribbean (Restrepo et al. 2018). The shallow portion of Varadero reef is in good condition in terms of coral cover (up to 50-60%) (Pizarro et al. 2017), despite its proximity to the Dique outlet and Cartagena city. A detailed description of the study sites is provided in the Supplementary Material.

**Fig. 1.**
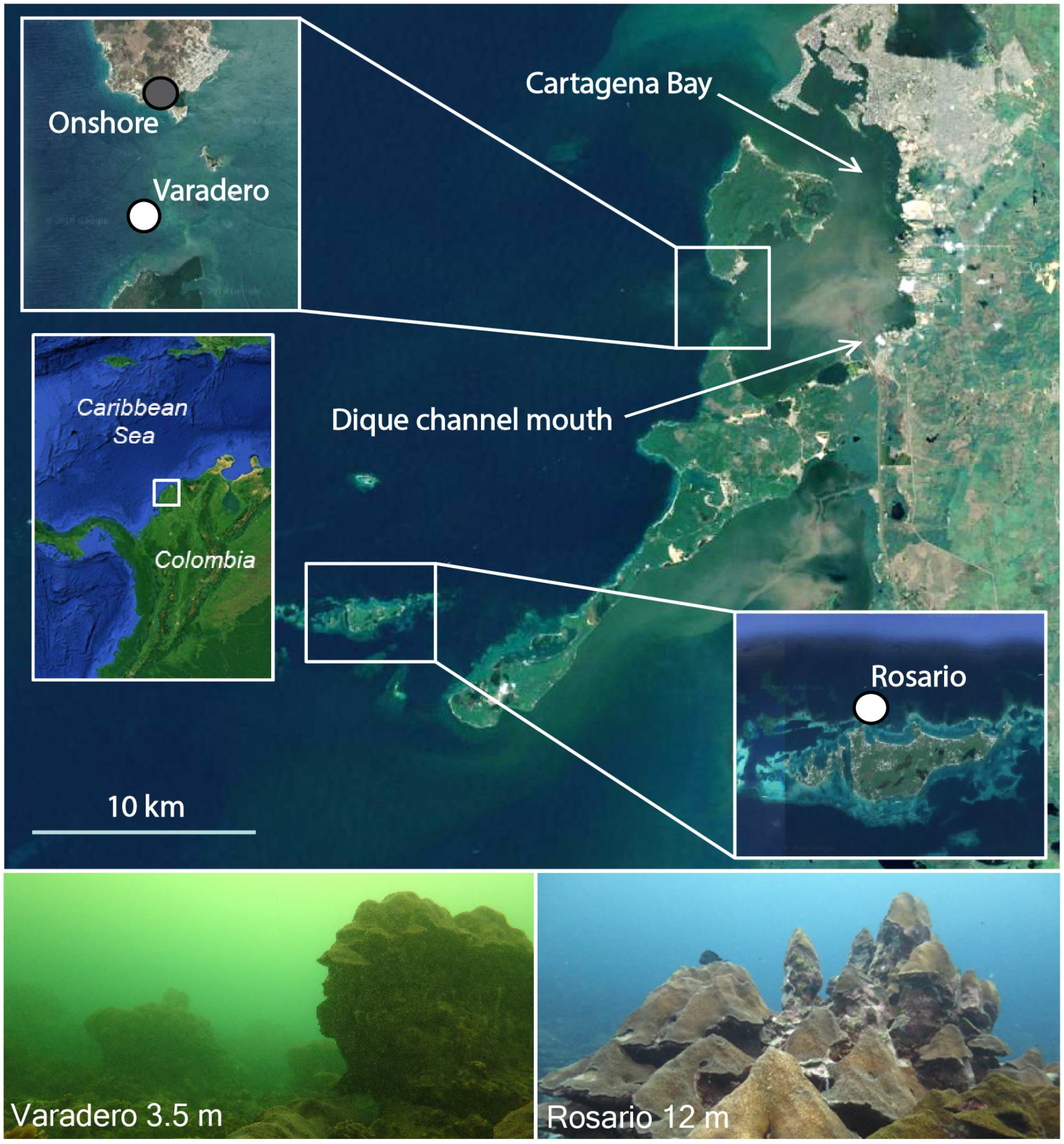
Study sites with contrasting exposure to the Dique plume. Varadero is located ~6 km west of Dique mouth, and Rosario 21 km southwest of Cartagena Bay (white circles). Surface irradiance was measured onshore close to Varadero (grey circle) to estimate *K*_d_ variation. Arrows indicate the location of Cartagena Bay and Dique channel mouth. Lower panels illustrate a general view of the study sites. Map data: Google, Maxar Technologies.

Preliminary analyses of the vertical diffuse attenuation coefficient for downwelling irradiance (*K*_d_) obtained by measuring light intensities across depths using the cosine-corrected PAR sensor of a Diving PAM (Walz, Germany), indicated that the total light exposure at 3.5 m in Varadero and at 12 m in Rosario was similar. Temperature at these depths was recorded every 30 min between November 2016 and July 2017 with HOBO pendant dataloggers (UA-002-64, Onset Computer Corporation, USA). A small (0.27 ºC) but significant difference in temperature was detected between sites (29.15 ± 1.22 (daily mean ± SD) in Varadero and 28.88 ± 0.89 °C in Rosario; *H*_(1)_ = 5.58, *p* = 0.018). However, the resulting variation in *O. faveolata* metabolic rates based on the scaling quotient of temperature (Q_10_) (Scheufen et al. 2017b) is estimated to be negligible (<4%).

### Coral Sampling

On October of 2016, coral fragments (10 cm^2^) were collected from the edge of 15 apparently healthy *Orbicella faveolata* colonies at each site. Source colonies were chosen randomly at a constant depth of ~3.5 m in Varadero and ~12 m in Rosario, where total light exposure was similar. Fragments were fixed to PVC panels with non-toxic epoxy (Z-Spar A-788 epoxy) placed at the depths of collection. Corals were allowed to recover and fully acclimate for seven months at each site. On May 2017, a subsample from the survivors (12 in Rosario and 15 in Varadero) was used for symbiont genotypification and physiological analysis.

### Irradiance measurements

Irradiance was monitored every ten minutes for one year (November 2016 - November 2017) at each site with cosine-corrected light sensors (Odyssey submersible PAR logger, Dataflow systems, New Zealand), previously cross-calibrated against a manufacturer-calibrated quantum sensor (LI-1400, LI-COR, USA). The light sensors were cleaned and downloaded periodically (every two months or less). Visual inspections at each visit showed that sediments and biofouling were not covering the sensors, potentially related to the type of sediments of the Dique plume (mainly fine silts and clays) which tend to remain suspended in the water column (Lonin et al. 2004). Even if the potential interference of sediments in the sensor’s signal is low, a linear regression on the data was used for correcting cumulative signal attenuation (Fig. S1).

### Spatiotemporal variation of water optical properties

To estimate the variation of the water optical properties resulting from the Dique plume dynamics, light data recorded underwater were compared with data simultaneously recorded onshore close to Varadero (Fig. 1). With this array, we isolated the variations of irradiance associated with the plume from variations due to cloud coverage. The irradiance synchronously recorded onshore and underwater was used to estimate instantaneous values of *K*_d_ based on the Lambert-Beer’s law:

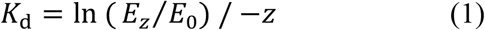

where *E*_z_ is underwater irradiance, *E*_0_ is onshore irradiance, and *z* is the depth of the underwater sensor (3.5 m). Only data recorded between 7:00 and 17:00 were employed. The analysis was performed over one-hour averages, smoothing out anomalies due to differences in cloud coverage between sites at short time intervals. Occasional failures of either sensor reduced the amount of time intervals that could be used to estimate *K*_d_’s, leaving a total of 86 days of usable data.

The vertical spectral diffuse attenuation coefficient for downwelling irradiance (*K*_d_ *λ*) was estimated by modifying the methodology of Maritorena and Guillocheau (1996). A mini-spectrophotometer (Flame T-UV-VIS-ES, Ocean Optics, USA) connected to a cosine-corrected sensor through a 30 m fiber optic cable (Avantes, Apeldoorn, The Netherlands) was used to measure in triplicate the irradiance at ocean surface and at several depths at each site. A bubble level was used to keep the light sensor horizontal, avoiding interference of divers or boat shading. *K*_d_ *λ* was calculated based on the change of underwater light spectra relative to surface. The spectral sensitivity of the instrument and the attenuation of the fiber optic were accounted for by normalizing the surface spectra against a traceable solar spectra reference (SORCE/SIM 2020).

### Symbiont identity

Coral samples were stored at -80 °C prior to processing. 50 μl of DNA was extracted from each sample (*n* = 15 in Varadero; *n* = 12 in Rosario) using the MoBio Powersoil DNA Isolation Kit (MoBio Laboratories, Carlsbad, California). Two-stage amplicon PCR was performed on the Internal Transcribed Spacer 2 (ITS2) rRNA marker gene. We used modified versions of the itsD and its2rev2 (Stat et al. 2009), including universal primer sequences that are required for Illumina MiSeq amplicon tagging and indexing. The PCR amplification was structured as follows: 2 min of denaturation at 94 °C; 35 cycles of 45 s at 94 °C, 60 s at 55 °C, and 90 s at 68 °C; then finally 7 min at 68 °C. Samples were sequenced using the Illumina MiSeq platform at the DNA Services Facility at the University of Chicago, Illinois. Resulting sequences were submitted to SymPortal for further processing and downstream analyses (Hume et al. 2019). Sequences within the Symbiodiniaceae family were identified and separated into genera, formerly “clades” (LaJeunesse et al. 2018), which were further separated into ITS2 type profiles representative of putative Symbiodiniaceae taxa (Hume et al. 2019). Sequences variants occurring in less than 1% of the sample were omitted in order to remove rare intragenomic variants and sequence artifacts generated by MiSeq sequencing.

### Photophysiology

Chlorophyll *a* (Chl *a*) fluorescence data were collected *in situ* with a submersible pulse amplitude modulated fluorometer (Diving-PAM, Walz, Germany). Details of PAM settings and environmental conditions during measurements are provided in the Supplementary Material. On cloudless days (*n* = 3 at each site), we quantified PSII photochemical yields in algal symbionts of 15 colonies randomly distributed near the experimental sites at constant depths (~3.5 m Varadero and ~12 m Rosario). Measurements were taken in the horizontal-uppermost part of colonies to avoid interference of intra-colony light gradients. The effective quantum yield (*ΔF/F*_m_*’*) of photosystem II (PSII) was recorded at local noon and the maximum quantum yield (*F*_v_*/F*_m_) of PSII at dusk or dawn. The maximum excitation pressure over PSII (*Q*_m_) was calculated following Iglesias-Prieto et al. (2004):

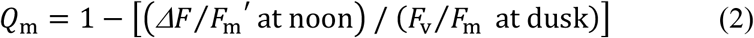

Chl *a* was extracted from tissue slurries (*n* = 7 Varadero; *n* = 8 Rosario) obtained with an air gun connected to a scuba tank and subsequently homogenized with a Tissue-Tearor Homogenizer (BioSpec Inc, USA). Pigment extraction was performed in acetone/dimethyl sulfoxide (95:5 vol/vol). Chl *a* density was estimated spectrophotometrically (*n* = 3 per sample) with a modular spectrometer (Flame-T-UV-VIS, Ocean Optics Inc., USA) using the equations of Jeffrey and Humphrey (1975). The specific absorption coefficient of Chl *a* (a^*^_Chl *a*_), descriptor of pigment light absorption efficiency of intact corals, was calculated according to Enríquez et al. (2005):

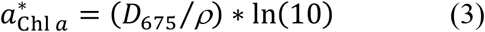

where *D*_675_ is the estimated absorbance value of corals at 675 nm, calculated from reflectance (*R*) measurements as [*D*_675_ = log (1/*R*_675_)], and *ρ* is the pigment content per projected area (mg Chl *a* m^-2^). Corals reflectance spectra was measured using a similar optical set-up as Vásquez-Elizondo et al. (2017). Coral surface area for these and other metrics was determined with the aluminum foil technique (Marsh 1970). Cell counts to estimate algal density and cell pigmentation were not reliable due improper sample preservation and not included in analysis.

Photosynthetic parameters were obtained from PE (photosynthesis vs irradiance) curves performed under laboratory conditions in a gradient of artificial light. Incubations (*n* = 8 in Varadero; *n* = 6 in Rosario) were performed in a custom-made acrylic chamber with four hermetic compartments (~650 ml each) filled with filtered seawater (0.45 µm) under constant agitation. Temperature was maintained at 28 °C with an external circulating water bath (Isotemp, Fisher Scientific). Ten levels of irradiance previously measured with a Diving-PAM light sensor were supplied at 10-min intervals with four 26 W LED bulbs (UL PAR38, LED Wholesalers Inc, USA) controlled with a custom-made software in continuous-mode. The range of light intensity covered was 0 to ~1400 µmol quanta m^-2^ s^-1^. Oxygen concentrations were measured with a 4-channel fiber-optic oxygen meter system (FireSting, Pyroscience). Parameters were calculated following Iglesias-Prieto and Trench (1994). Additional information on the light system and photosynthetic parameters is provided in the Supplementary Material.

### Ecological survey

We performed ecological surveys to estimate the condition of *O. faveolata* populations. Random dives near each site allowed us to define the vertical distribution ranges of the species. Colony abundance and percentage of old mortality were recorded across five sequential 10×1 m belt transects deployed perpendicular to the reef slope at six depths in Varadero (2, 3, 4.5, 6, 7.5 and 9 m) and Rosario (3, 5, 7, 9, 13 and 17 m). The percentage of old mortality was estimated following the AGRAA protocol (*i*.*e*., the skeleton is no longer white and has been lost or covered by epibenthic organisms) (Lang 2003). Completely dead colonies were excluded from surveys due to the typical loss of corallite skeletal structures that allow to differentiate species.

### Integrated productivity

Variation of the phototropic contribution of algal symbionts to the energy requirements of *O. faveolata* colonies across depths was estimated based on daily integrated photosynthesis to respiration [P:R] ratios. P:R ratios were estimated using the PE curve parameters (mean values) and light availability across depths, calculated from the estimated *K*_d_’s (derived from equation 1). Daily integrated photosynthesis 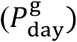 was calculated with a hyperbolic tangent function (Jassby and Platt 1976):

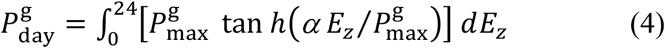

where 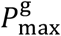is maximum gross photosynthesis, *α* is photosynthetic efficiency, and *E*_*z*_ is the estimated irradiance at depth *z*. The daily integrated respiration (*R*_day_) was calculated with a hyperbolic tangent function modified from Chalker (1981):

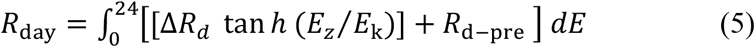

where Δ*R*_d_ represents the difference between the post- and pre-illumination respiration rates, *E*_*k*_ is saturating irradiance, and *R*_d*-*pre_ is pre-illumination respiration rate. We assumed a light-associated asymptotic increase of the respiratory activity until reaching a maximum determined by *E*_k_. This assumption is based on evidence that indicate that respiration rates in corals are light-driven and closely coupled with the internal oxygen concentration (Colombo-Pallotta et al. 2010; Holcomb et al. 2014) (Fig. S2).

## Results

### Spatiotemporal variation of water optical properties

Analysis of *K*_d_’s obtained at the beginning of the study with a Diving-PAM light sensor indicated a strong stratification of the water column in Varadero (Fig. 2a). The stratification was characterized by a superficial layer of ~1 m with an extremely high *K*_d_ (1.93 ± 0.26 m^-1^, mean ± SD), and a clear subsurface layer with significantly lower *K*_d_ (0.31 ± 0.11 m^-1^) (*t*_(7)_ = 15.31, *p* < 0.01). No noticeable stratification was detected in Rosario, which revealed a monotonic *K*_d_ (0.16 ± 0.01 m^-2^) (Fig. 2a).

**Fig. 2.**
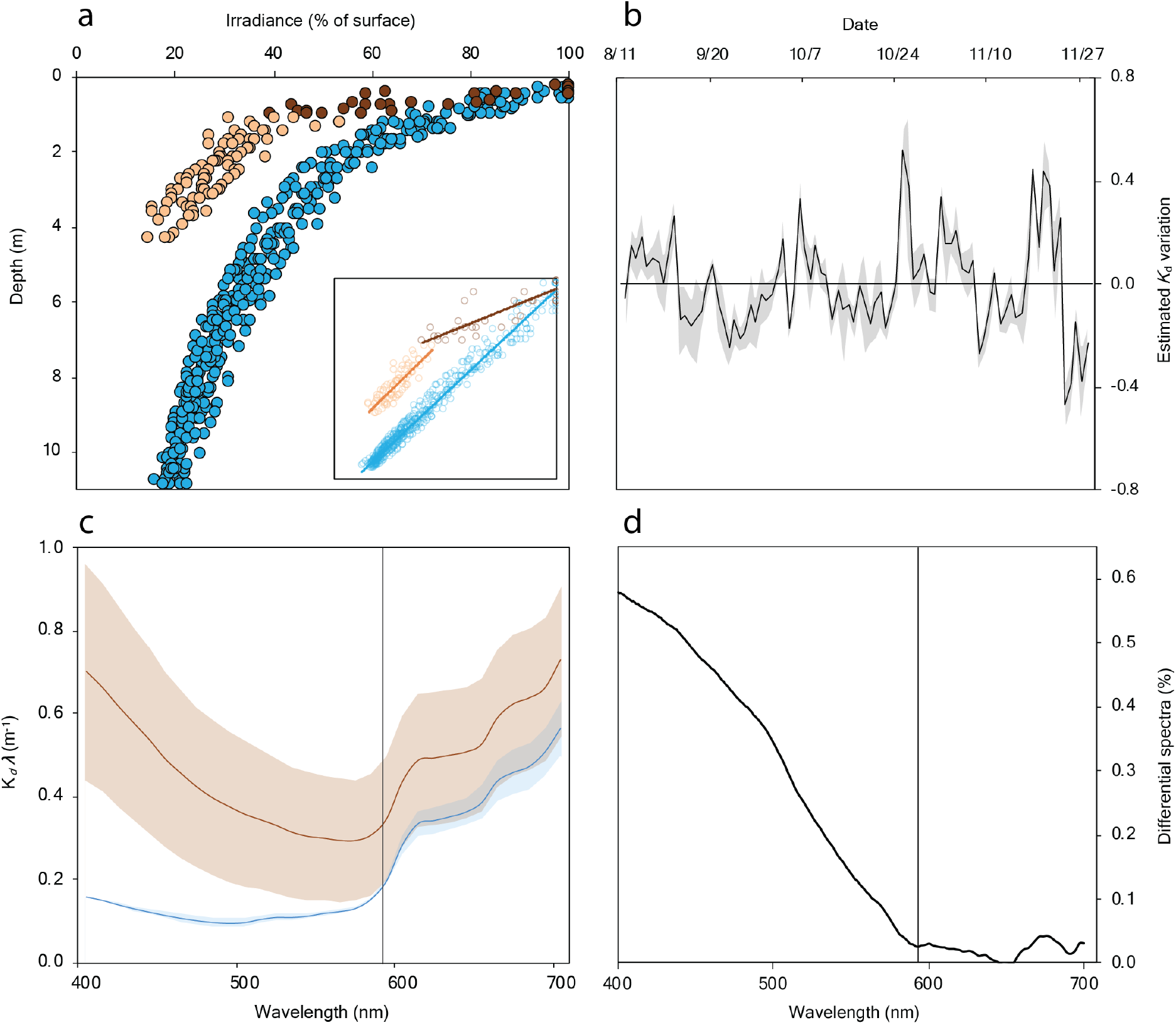
Variation of underwater light climate and water optical properties. **a** Strong stratification of the water column in Varadero characterized by a superficial layer with high *K*_d_ (1.93 ± 0.26 m^-1^, dark brown) and a clearer subsurface layer with low *K*_d_ (0.31 ± 0.11 m^-1^, light brown), compared with Rosario monotonic *K*_d_ (0.16 ± 0.01 m^-1^, blue). The insert shows light data distribution with depth in log-scale. **b** Temporal variation of mean daily *K*_d_ in Varadero estimated from onshore and underwater irradiances (shaded area represents SD). **c** *K*_d_ *λ* in Varadero (brown) and Rosario (blue) (shaded areas represent 95% confidence intervals). **d** Estimated light attenuation due to absorptance of dissolved organic matter associated to Dique discharges based on a differential normalized spectrum in Varadero. Vertical line in **c** and **d** separate the two contrasting spectral regions.

Estimated *K*_d_’s from the synchronous oscillation of onshore and underwater irradiances evidenced a strong oscillation of water optical properties in Varadero (Fig. 2b). Estimated *K*_d_’s were 0.92 ± 0.26 m^-1^ and ranged between 0.38 and 1.95 m^-1^. Extremes within this range indicate that underwater irradiance can vary by up to two orders of magnitude only due to changes in water optical properties. Maximum *K*_d_’s were higher compared to previous reports at this site (Pizarro et al. 2017; Roitman et al. 2020) and other turbid-water reefs (Hennige et al. 2010; Omachi et al. 2019).

Analyses of *K*_d_ *λ* indicate distinctive light scattering and absorbing characteristics of dissolved and particulate matter in the water at each site (Fig. 2c). Varadero’s high variability of *K*_d_ *λ* (*i*.*e*., wide confidence intervals) (Fig. 2c) is consistent with the strong stratification (Fig. 1a) and the temporal variation of *K*_d_ (Fig. 2b). *K*_d_ *λ* evidenced two spectral regions with contrasting characteristics. One below ~600 nm with increasing attenuation toward shorter wavelengths in Varadero and low attenuation in Rosario, attributed to light absorption of dissolved organic material of continental origin (Maritorena and Guillocheau 1996). And a spectral region above 600 nm with nearly constant properties despite differences in attenuation magnitude among sites, attributed to light attenuation due to scattering by particulate matter with small effect on spectral quality (Fig. 2c). The estimated attenuation due to absorptance of dissolved organic matter obtained from a differential normalized spectrum (Fig. 2d), demonstrates the high concentration of dissolved organic matter in Varadero associated with the Dique plume and its critical role in modifying the light quality.

Dramatic changes of *K*_d_ were detected within and between days (Fig 3), highlighting the variable nature of the water optical properties and underwater light climate associated with the Dique plume dynamics in Varadero. On average, the daily integrated irradiance was 7% lower in Varadero than in Rosario (2.03 ± 1.32 and 2.17 ± 0.91 mol quanta m^-2^ d^-1^, respectively) (*H*_(1)_ = 7.92, *p* < 0.05). Prevalent irradiance at both sites was consistent with low-light reef environments (Anthony and Hoegh-Guldberg 2003; DiPerna et al. 2018).

**Fig. 3.**
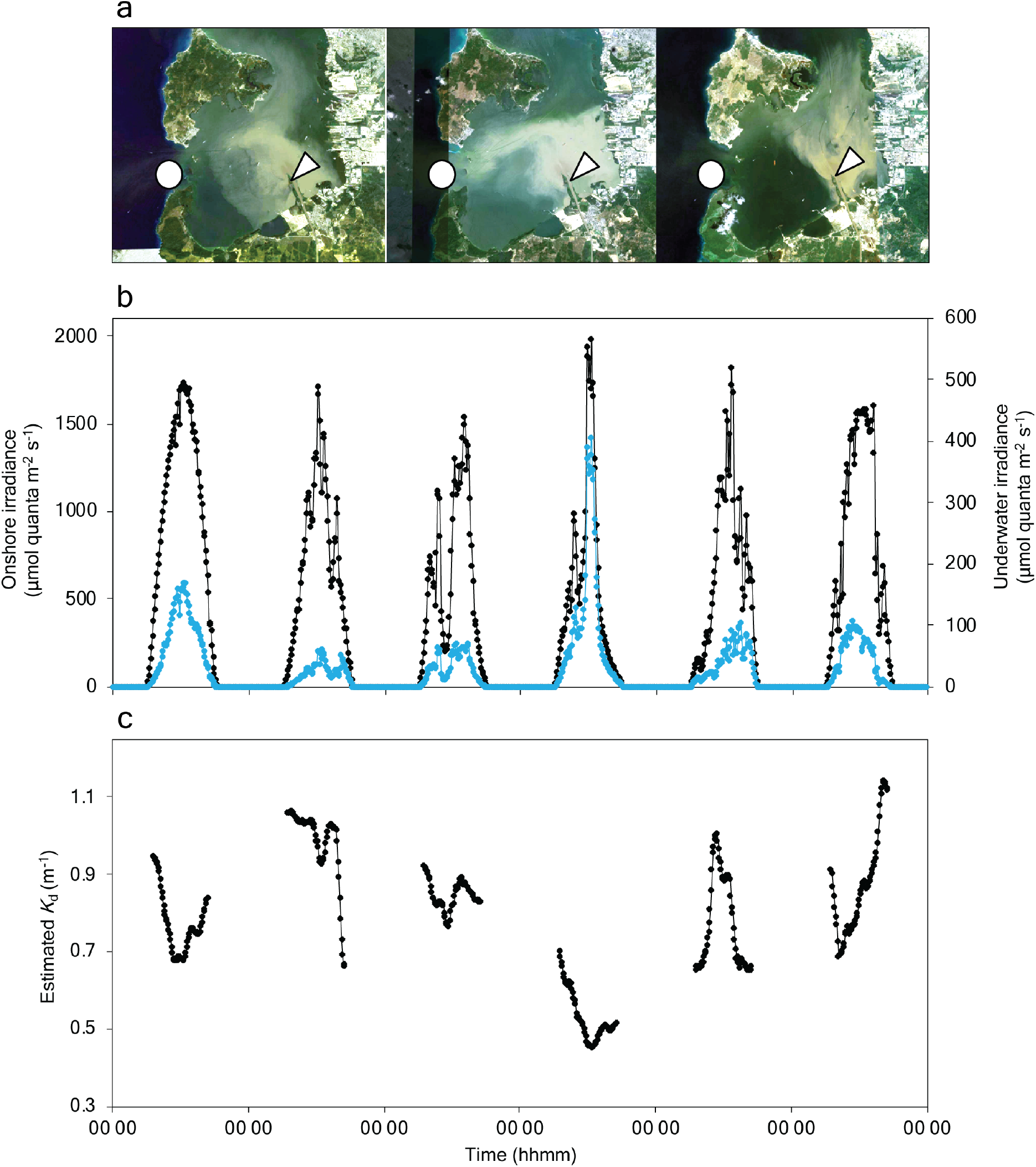
Effect of Dique plume dynamics on *K*_d_. **a** Satellite images illustrating contrasting spatial patterns of the plume, signaling the location of Varadero (circle) and the Dique mouth (arrow). **b** Oscillation patterns of onshore (black line) and underwater (blue line) irradiances on six random days. **c** *K*_d_ variation estimated from onshore and underwater irradiances.

### Symbiont identity

Analyses of Internal Transcribed Spacer 2 (ITS2) rRNA sequences identified distinctive ITS2 profiles combinations diagnostic of five separate genotypes. One belonged to the genus *Symbiodinium* (ITS2 type A3), and three to *Cladocopium* (ITS2 types C7, C7/C12c, C3ee/C21/C3an) (Fig. 4, Fig. S3). A *Breviolum* genotype (ITS2 type B1) was only detected in some corals from Rosario but in low background levels. *Symbiodinium* ITS2 type A3 occurred in colonies from both locations and in some samples, it was the only dominant symbiont. Many colonies from Varadero contained a *Cladocopium* genotype (ITS2 type C3ee/C21/C3an), which occurred alone or in combination with *Symbiodinium* ITS2 type A3. This *Cladocopium* genotype was also found in a few corals from Rosario as the dominant symbiont. Two additional albeit related *Cladocopium* genotypes (ITS2 types C7 and C7/C12c) were also present in certain colonies from Rosario, but never together. Statistical tests confirmed that the Symbiodiniaceae communities in terms of ITS2 profiles composition were different between sites (*p* = 0.001, PERMANOVA on Bray-Curtis matrix).

**Fig. 4.**
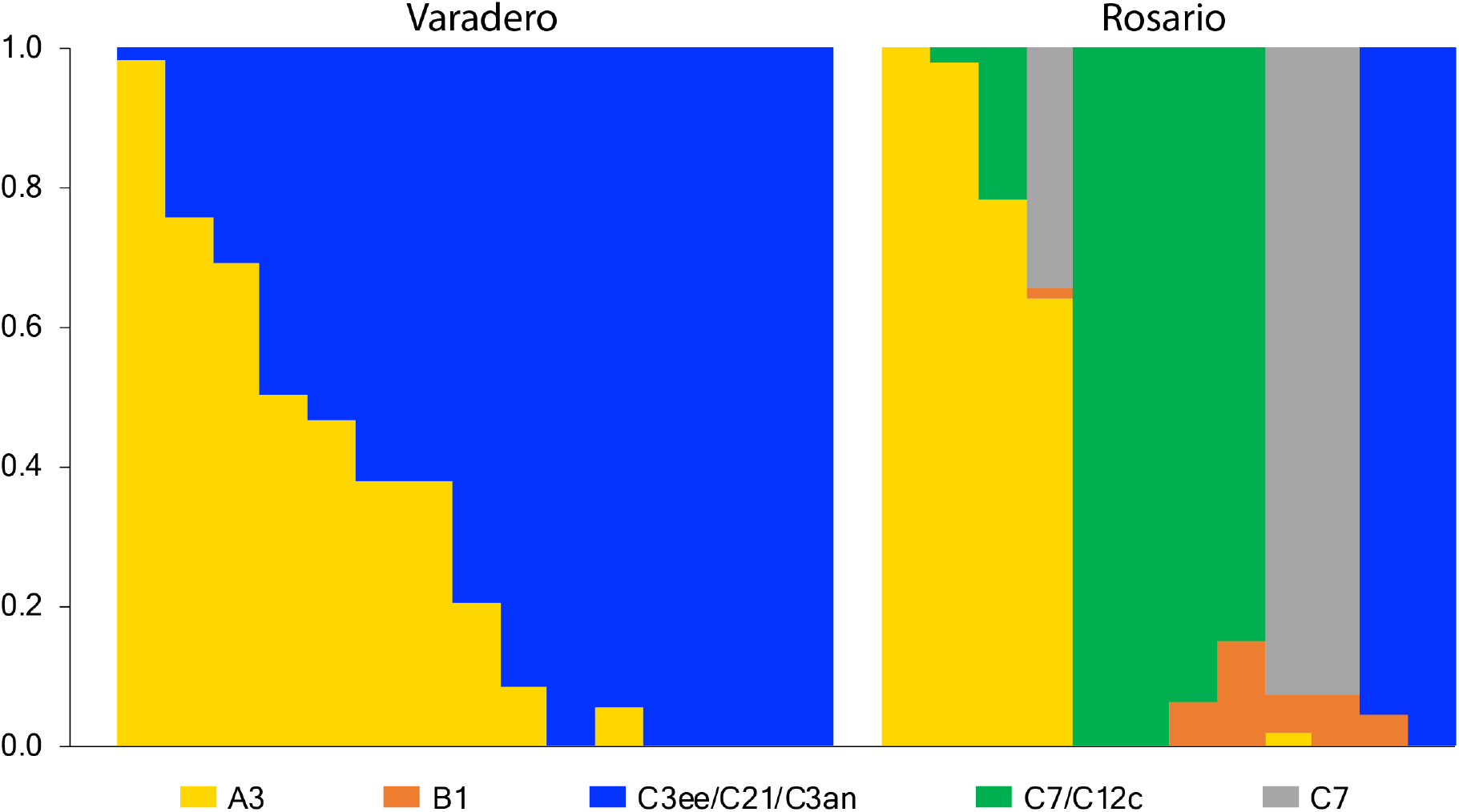
Predominant Symbiodiniaceae ITS2 profiles associated to corals from Varadero and Rosario. Each column of stacked bars represents one sample, with bar heights within each column representing the relative abundance of sequences. Raw diversity of ITS2 type profiles is provided in Fig. S3.

### Photophysiology

Photosynthetic parameters derived from the PSII photochemical yield of symbionts (*F*_v_*/F*_m_, *ΔF/F*_m_*’* and *Q*_m_) and host pigmentation (Chl *a* content and a*_Chl *a*_) were significantly different between sites. Parameters derived from the photosynthetic potential (*α, E*_c_, *E*_k_, *R*_d_, *P*_max_ and 1/*Φ*_max_) and corals capacity to absorb light (A_PAR_) were non-significantly different (Table 1). *F*_v_*/F*_m_ values (0.669 ± 0.019 and 0.676 ± 0.012 in Varadero and Rosario, respectively) differed by 1% (*t*_(74)_ = 2.26, *p* = 0.027). Values of *ΔF/F*_m_’ and *Q*_m_ (0.579 ± 0.052 and 0.610 ± 0.033, and 0.135 ± 0.062 and 0.097 ± 0.051 in Varadero and Rosario, respectively), varied by 5% (*t*_(75)_ = 3.38, *p* = 0.001) and 28% (*t*_(84)_ = -3.09, *p* = 0.003), respectively. Estimated values of 1/*Φ*_max_ (11.24 ± 1.87 and 11.62 ± 3.16 quanta O_2_^-1^ in Varadero and Rosario, respectively) were lower than previous reports (Wyman et al. 1987; Rodriguez-Roman et al. 2006) and close to minimum practical limits of ~10-12 quanta O_2_^-1^ (Hill and Govindjee 2014).

**Table 1.**
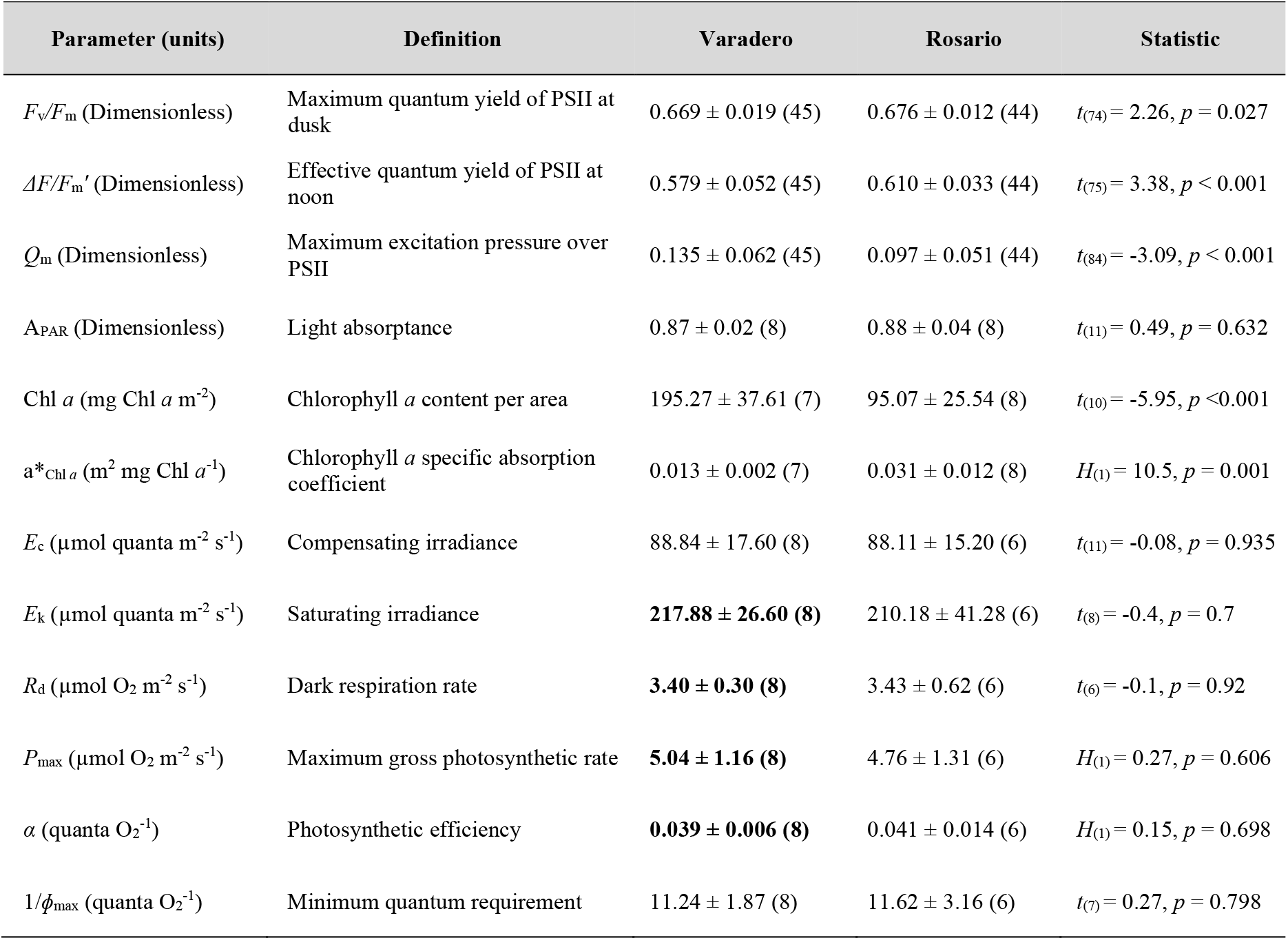
Photoacclimation parameters of *O. faveolata* in Varadero at 3.5 m and Rosario at 12 m. Values correspond to the mean ± SD (sample size *n*). *t*-tests (*t*) and Kruskal-Wallis tests (*H*) were conducted after testing of assumptions (normality with the use of K-S test) to determine if differences between sites were statistically significant. Values of photosynthetic parameters used to estimate the productivity of colonies (integrated daily P:R ratios) are highlighted in bold.

Chl *a* density was significantly higher (*t*_(10)_ = -5.95, *p* < 0.001) in corals from Varadero compared to Rosario (195.27 ± 37.61 and 95.07 ± 25.54 mg Chl *a* m^-2^, respectively). Changes in Chl *a* density resulted in significant variations of a*_Chl *a*_ (0.013 ± 0.002 and 0.031 ± 0.012 m^2^ mg Chl *a*^-1^ in Varadero and Rosario respectively, *H*_(1)_ = 10.5, *p* = 0.001). Comparative analyses of changes in a*_Chl *a*_ as a function of Chl *a* content indicate a relatively steady light harvesting efficiency despite larger differences in pigment content in corals from Varadero. In contrast, reduced Chl *a* content variation in Rosario resulted in more dramatic changes of the effective absorption cross section of algal pigments (Fig. S4). Our data provide no clear evidence that these differences were associated with distinctive Symbiodiniaceae composition (Fig. S5). No significant differences were found in light absorption capacity (A_PAR_) and photosynthetic descriptors (*α, E*_c_, *E*_k_, *R*_d_ and *P*_max_) between sites (Table 1).

### Ecological survey

*O. faveolata* vertical distribution ranged between 2 and 9 m in Varadero, and 3 and 17 m in Rosario. Varadero had both higher density of colonies (3.20 ± 2.85 colonies transect^-1^) and coral cover (4.46 ± 4.57 m^2^ transect^-1^) than Rosario (1.70 ± 1.18 colonies transect^-1^ and 4.20 ± 5.48 m^2^ transect^-1^, respectively). The highest abundance (38% of total colonies) occurred at 4.5 m in Varadero and at 9 m (33%) in Rosario (Fig. S6). In Varadero, the lowest partial colony mortality occurred at 2 m (7.5 ± 3.5%) and the highest at 9 m (73.3 ± 26.4%). In Rosario, the lowest mortality occurred at the maximum depth (5.0 ± 12.2% at 17 m) and the highest at an intermediate depth (73.3 ± 20.8% at 5 m) (Fig. 5). Depth and mortality had a strong exponential relationship in Varadero (*R*^2^ = 0.77, *p*<0.01) and a non-significant linear (*R*^2^ = 0.43, *p* = 0.16) or exponential (*R*^2^ = 0.33, *p* = 0.23) relationship in Rosario.

**Fig. 5.**
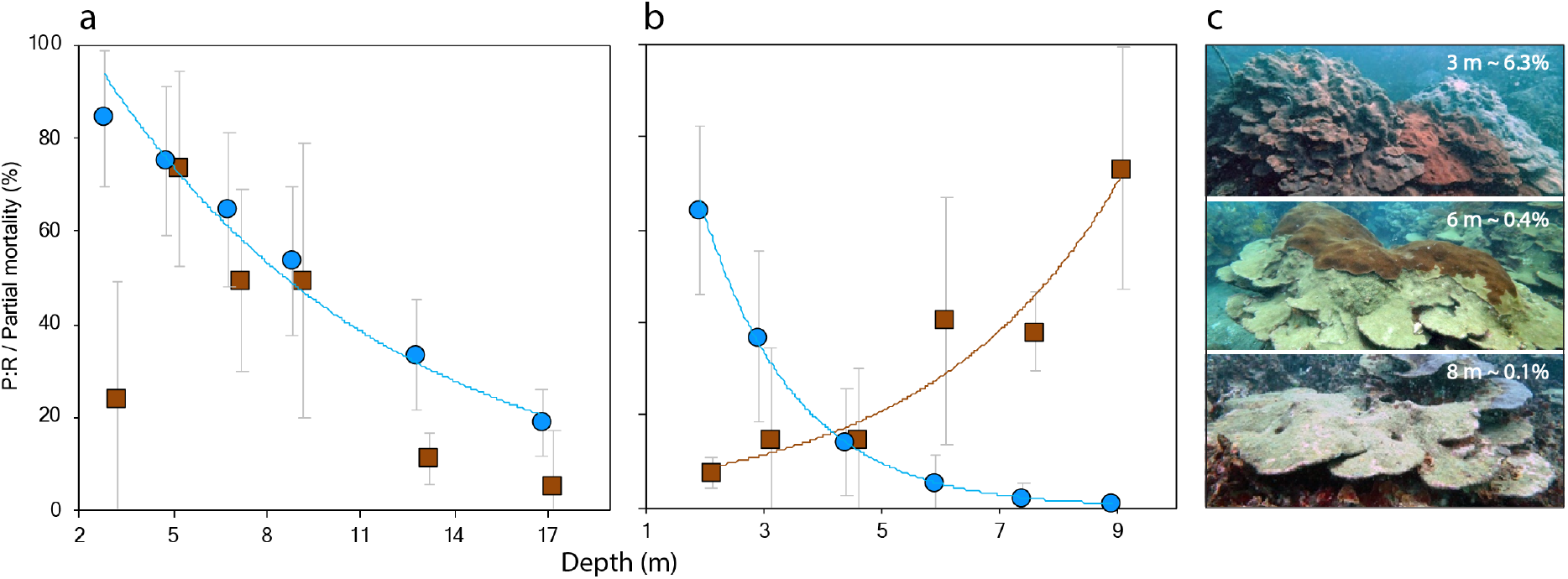
Productivity and mortality of *O. faveolata* across depths. Partial mortality (brown squares) and daily integrated P:R ratios (%, blue circles) in Rosario (**a**) and Varadero (**b**). Values correspond to mean ± SD. Exponential regressions were used to fit the data (only significant relationships are depicted). **c** Mortality patterns across depths in Varadero, indicating the % surface light at each depth based on the mean estimated *K*_d_.

### Integrated productivity

Integrated daily P:R ratios of *O. faveolata* colonies in Varadero suggest a variable phototrophic contribution of algal symbionts to the metabolic demand of coral holobionts, both spatially (across depths) and temporally, as a consequence of changes in light availability. At 2 m depth, it is estimated that symbionts can supply, on average, 64% of daily holobiont’s energy requirements (P:R = 0.64 ± 0.18). On days with increased light penetration the symbionts supply 99% of holobiont’s requirements while on days with highest *K*_d_’s the supply drops to ~20%. At 9 m, the maximum depth where living *O. faveolata* colonies are currently found in Varadero, the contribution of symbionts is, on average, 1% of holobiont’s energy requirements, although it drops to practically zero during days with highest *K*_d_’s (P:R = 4.49 × 10^−6^) (Fig. 5). A strong, negative correlation was found between colonies partial mortality and daily P:R ratios (Spearman *r*_s_ = -0.94, *p* = 0.005). The coefficient of determination of an exponential regression indicates that the phototrophic contribution explains 78% of the mortality variation across depths in *O. faveolata* (Fig. S7).

In Rosario at 3 m, the shallowest depth of *O. faveolata*, it is estimated that the symbionts supply, on average, 84% of the holobiont’s energy requirements (P:R = 0.84 ± 0.14). At 17 m, the maximum depth of *O. faveolata*, the symbionts supply is of 19% (P:R = 0.19 ± 0.07), whit minimum and maximum values of 2% and 27% depending on changes in light availability (Fig. 5). Rosario had significantly higher estimated P:R ratios than Varadero both at the upper (*H*_(1)_ = 73.29, *p* < 0.001) and lower (*H*_(1)_ = 189.25, *p* < 0.001) local depth limits of *O. faveolata*. A non-significant correlation was found between colonies partial mortality and estimated P:R ratios in Rosario (Spearman *r*_s_ = 0.60, *p* = 0.208).

## Discussion

We identified three main features of the underwater light climate in Varadero, a coastal reef exposed to a turbid plume of water originating from a man-made channel: 1) light-limiting conditions due to the strong attenuation generated by particulate matter and dissolved substances concentrated in a superficial layer, 2) substantial variation of the underwater light climate in response to fluctuations on water optical properties, and 3) altered spectral composition of light due to the wavelength-selective absorption by organic matter of continental origin (Maritorena and Guillocheau 1996; Hennige et al. 2010). These patterns are produced by the freshwater discharges with high sediment load from the Dique channel in the vicinity of the reef. The strong stratification, also reported in previous studies (Pizarro et al. 2017; Tosic et al. 2019), results from the different properties (*e*.*g*., salinity and temperature) of a superficial layer directly affected by the Dique channel discharges and a subsurface clear layer of oceanic waters (Lonin et al. 2004). The strong stratification of the water column seems to isolate the Varadero reef from the Dique freshwater discharges but have a critical impact on the underwater light availability and photosynthetic energy acquisition in corals.

The variation of water optical properties in Varadero explains 81% of the overall temporal variation of underwater irradiance. This estimation is in close proximity to irradiance variation in other turbid-water reefs due to suspended solids (Anthony et al. 2004). The observed variability is attributed to meteorological conditions affecting the Dique plume dynamics (Lonin et al. 2004; Tosic et al. 2019). At Rosario, although the water column is significantly clearer than Varadero, the *K*_d_’s for the whole PAR range (0.158 m^-1^) and towards the blue part of the spectrum (0.160 m^-1^ at 400 nm) were still notably higher compared to other reports in clear waters (Maritorena and Guillocheau 1996). This suggests an increased attenuation likely associated with the Dique discharges, highlighting the strong influence of continental runoff on the underwater light climate far beyond the channel mouth.

Our results suggest that *O. faveolata*, a major reef-building coral in the Caribbean, hosts distinctive Symbiodiniaceae communities in Varadero and Rosario. Two features of the Symbiodiniaceae communities are highlighted: 1) Despite the extreme fluctuations of the water optical conditions in Varadero, *Durusdinium trenchii* (ITS2 type D1a) was not detected in coral samples at this site. This known stress-tolerant species is adapted to survive in regions exposed to high temperature and irradiance (LaJeunesse et al. 2009). Given the distinctive features of Varadero reef environment, *D. trenchii* would be expected to thrive in *O. faveolata* colonies, but instead *Cladocopium* ITS2 type C3 and *Symbiodinium* ITS2 type A3 were dominant. And 2) corals from Rosario showed a greater diversity of Symbiodiniaceae communities, mainly due to an increased richness of *Cladocopium* genotypes. The reduced diversity of the symbiont community in corals from Varadero, similar to other turbid marginal reef environments (Smith et al. 2020), may be an indicator of the strong selective pressure exerted by the turbid plume dynamics on the symbiont communities.

*O. faveolata* corals display similar phenotypes in Varadero at 3.5m and Rosario at 12m. This highlights the essential role of the water optical properties on the underwater light climate and illustrates the constraint of using depth as main proxy for corals photoacclimation status. Estimated minimum quantum requirements (1/*Φ*_max_) of corals growing at both sites indicate similar light utilization efficiency close to theoretical operational limits of ~10-12 quanta O_2_^-1^ (Hill and Govindjee 2014). *Q*_m_ values close to 0 indicate that even at noon when corals are exposed to maximal irradiances, most PSII reaction centers of symbiotic algae remained open suggesting light-limited photosynthetic rates (Iglesias-Prieto et al. 2004). Both *Q*_m_ and 1/*Φ*_max_ normally decrease with increasing depth until reaching the minimal practical values, which corresponds to the limits of potential photoacclimation capacity and tolerance range for the symbiosis (Wyman et al. 1987; Iglesias-Prieto et al. 2004). The values of *Q*_m_ and 1/*Φ*_max_ close to theoretical minimums suggest that the photoacclimation of *O. faveolata* symbionts is close to the limit for maximum efficiency of solar energy utilization and, therefore, the lower vertical distribution limit of the species at each site despite the contrasting depth differences.

Although the average daily irradiance at both sites is consistent with low-light environments (Anthony and Hoegh-Guldberg 2003; DiPerna et al. 2018), corals from Varadero are occasionally exposed (~10% of days) to supersaturating irradiances that exceed the saturation point, *E*_k_, of algal symbionts. Irradiances at local noon during these days can be almost twice the estimated *E*_k_, representing a potential source of over-excitation of the photosynthetic apparatus that must be dissipated by photoprotective mechanisms. A greater reduction of the quantum yield of photosystem II (PSII) measured at noon (*ΔF/F*_m_*’*) relative to its maximum value at dusk (*F*_v_/*F*_m_), indicate that corals at Varadero were indeed exposed to higher peaks of irradiance during these measurements, compared to Rosario. It must be stressed, however, that the almost identical values of *F*_v_*/F*_m_ between sites indicate similar photochemical energy conversion efficiency of the PSII of coral symbionts. This suggests that the occasional exposure to higher irradiances in Varadero are not intense enough as to induce a chronic photoinactivation of a population of PSII reaction centers in order to increase the capacity to dissipate excess excitation via non-photochemical quenching (Hoegh-Guldberg and Jones 1999). Consistently, comparative analyses of the parameters derived from the P-E curves indicate that corals at both locations have very similar photosynthetic potential. Collectively, these results indicate that the occasional exposure to supersaturating irradiances in Varadero does not compromise the photosynthetic performance of symbionts.

Interestingly, corals in Varadero had greater Chl *a* content than in Rosario, which resulted in a significant reduction of the light absorption efficiency of algal pigments, a*_Chl *a*_, due to pigment self-shading (Scheufen et al. 2017a). Differences in a*_Chl *a*_ indicate that Varadero highly pigmented corals are less efficient at collecting light per unit of pigment. Although a detailed analysis of the water quality in Varadero is beyond the scope of this study, we speculate that the nutrient enrichment that exists within Cartagena Bay due to the Dique discharges (Tosic et al. 2019) may also extend to Varadero. It is known that nutrient enrichment interference on coral’s metabolism can result in increased algal densities and/or Chl *a* content within cells (Shantz and Burkepile 2014), which in turn can affect the photosynthetic potential of the coral. The nearly constant capacity of corals to absorb ~90% of incident light at both sites, despite differences in Chl *a* content and a*_Chl *a*_, indicate that the increased pigmentation in corals from Varadero does not confer them with any additional advantage in terms of light harvesting. This condition may be related with the particular capacity of *O. faveolata* to maximize light absorption at low pigmentation (Scheufen et al. 2017a). The lack of differences among photosynthetic parameters indicates that the light absorbed is also utilized with similar efficiency in photosynthesis by corals at both sites, and that a potential nutrient enrichment linked to Dique discharges is not affecting the photosynthetic potential of *O. faveolata*. Although it is known that the symbiont genotype can influence the coral host physiology (Kemp et al. 2014), we found no evidence that the differences in pigmentation and efficiency to collect light were connected with distinctive symbiont genotypes.

The productivity model based on P:R ratios reveals that the daily phototrophic contribution of algal symbionts is highly variable. The reduction of primary production with increasing depth and during exposure to periods of high light extinction could be counterbalanced by photoacclimation and heterotrophic plasticity (Anthony and Fabricius 2000). The low-light phenotypes with physiological metrics signaling maximal theoretical limits of photoacclimation in Varadero suggests that enhance feeding on suspended particles originating from the plume may be the more relevant alternative to compensate for reduced photosynthesis. However, the Dique plume particles not only represent a potential source of nutrition but also a stress factor for coral colonies with even lethal effects due to the energetically costly process of particle clearance and microbe-mediated infections favored by sedimentation (Junjie et al. 2014; Pollock et al. 2014). The minimum irradiance at which photosynthesis outweighs respiration (*E*_c_) is estimated to be exceeded only above 6 m, at least for short periods of time. Corals clearance capacity below this depth is expected to be strongly limited by the negligible phototrophic contribution of algal symbionts and the metabolic depression of the holobiont. Due to photosynthetic energy deprivation and physical disturbance of particulate matter, coral colonies would not be expected to survive at much greater depths.

Colonies censused in this study were sexually mature (> 100 cm^2^) and some were probably hundreds of years old based on their height (>2 m), which suggests that the local population of *O. faveolata* was established in Varadero before the Dique major rectification works (Mogollón 2013). Current vertical distribution is restricted to a much narrower and shallower depth range (2-9 m) compared not only with Rosario (3-17 m) but also with other clear-water sites in the Caribbean (*e*.*g*., 2-20 m in Belize (Pandolfi and Budd 2008) and 3-25 m in Curacao (Van Veghel 1994)). The compressed vertical distribution, together with the strong correlation between partial colony mortality and depth, suggest the occurrence of a progressive decline of the coral population from the bottom to the top of the reef. Although other environmental impacts linked to the plume that were not accounted for in this study may also lead to corals mortality, the physiological evidence suggest that the current condition of *O. faveolata* population in Varadero is associated with light-limiting conditions and photosynthetic energy deprivation as a result of the progressive degradation of water optical properties (Fig. 6).

**Fig. 6.**
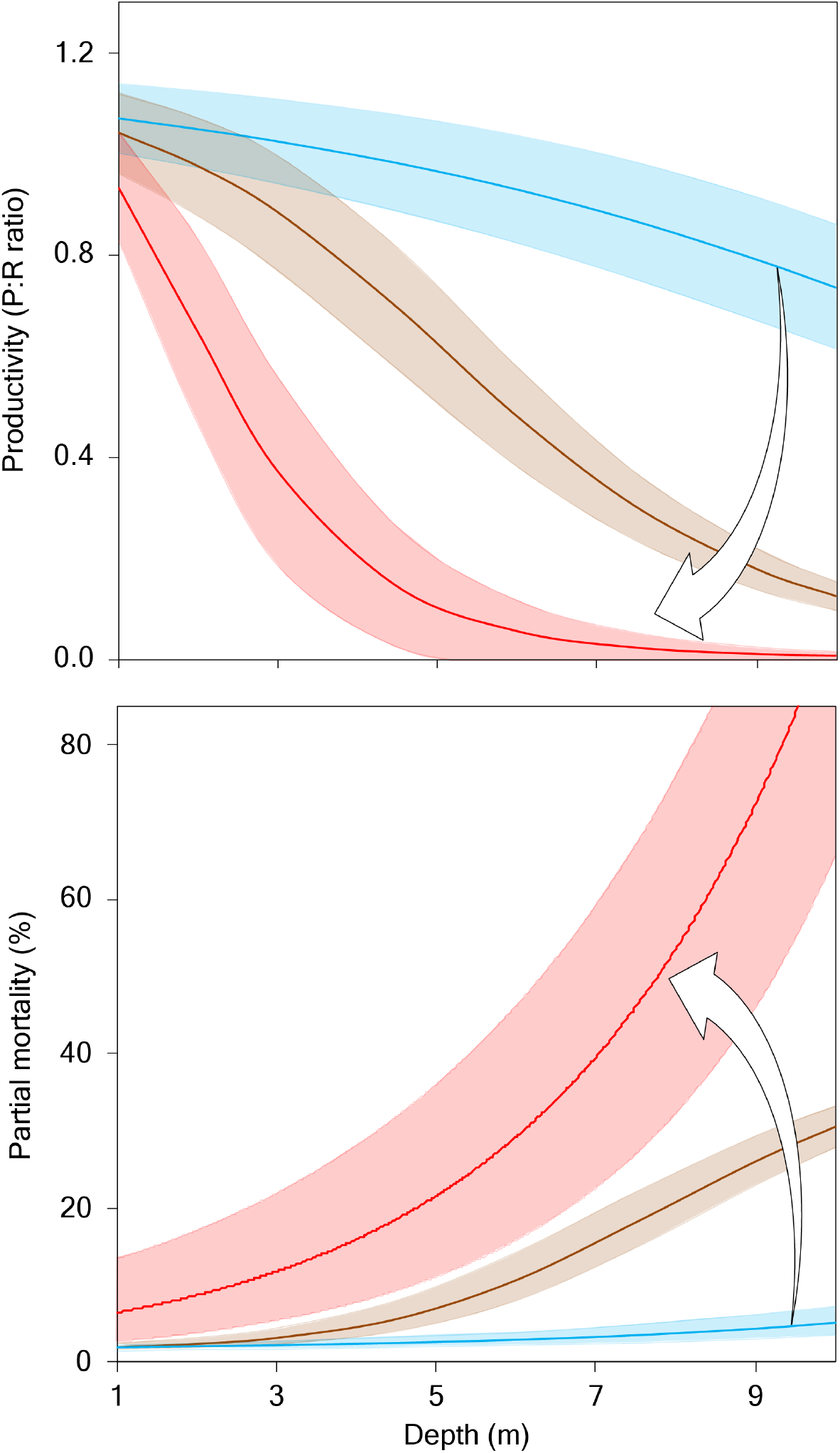
Predicted effect of the progressive degradation of water optical properties on colonies productivity and partial mortality across depths. Top panel: Estimated productivity (daily P:R ratios) with water of different *K*_d_’s. Lower panel: Predicted partial colony mortality based on the exponential regression describing the relation between P:R ratios and mortality. Responses were modelled under current *K*_d_ variation (red); under persistent low-intensity effect of the Dique plume (lowest estimated *K*_d_, 0.34 m^-1^) (brown), and under minimal influence of the plume (*K*_d_ of Rosario, 0.16 m^-1^) (blue).

Recent studies have documented higher coral cover and/or lower incidence of coral bleaching in turbid reefs potentially connected with an alleviating effect of reduced light levels in the water column (van Woesik et al. 2012; Morgan et al. 2017). This evidence has supported the idea that marginal coral reefs may act as refuges during the current coral reef crises. Our results partially support this hypothesis in the sense that a potential stress-relieving effect linked to turbidity could help explain the reduced partial mortality of *O. faveolata* colonies only in the shallowest portion of Varadero (< 4.5 m). However, with increasing depth, the elevated mortality and total absence of living colonies of *O. faveolata* below 9 m indicate that the gains from a potential alleviating effect are exceeded by the losses associated with the photosynthetic energy depletion resulting from light limitation.

Most coral reefs will be lost in a few decades unless there is a major scaling-up of management efforts and commitment based on an improved understanding of current ecological processes (Kennedy et al. 2013). Despite the significant steps taken to conserve coral reefs and their ecosystem services, the success of management efforts at large scales is low. In addition, major threats, such as global warming and ocean acidification, keep rising and driving reefs toward the breaking point of collapse (Ainsworth et al. 2016). Our findings suggest that effective policies seeking to improve water optical properties at local and regional scales should be considered a priority goal for coral reef conservation, a challenge that extends far beyond the limits of the coastal zone environment. If no action is taken to control the water optical quality and the current trajectory of increasing the flux of sediments into the coastal zone persists (Burke et al. 2011; Carlson et al. 2019), populations of reef-building corals like *O. faveolata* will be increasingly restricted to shallower depths, thereby eroding the ecosystemic benefits that these species provide.

## Acknowledgments

This study was funded by NSF grants OCE 1642311 and OCE 1442206 to MM and RIP, and Pennsylvania State University SSRI and IEE grants. B. Hume, T. LaJeunesse, and C. Prada provided help on Symbiodiniaceae analysis. M.F. Porras and D. Brown provided comments on data analysis. G. Navas, A. Bermúdez and D. Mendez from Universidad de Cartagena; J. Rojas and R. Vieira from Oceanario-CEINER; A.A. Shantz from Penn State University; the dive shops Cartagena Divers and Scuba Cartagena; and the Avendaño family provided extensive logistical support during field activities. The light system for PE curves was constructed by M.A. Gómez-Reali from Universidad Nacional Autónoma de México. E. Zarza from the Corales del Rosario y San Bernardo National Natural Park provided administrative and logistical support. The research was conducted under the collection permit No. 0546 from 2014 issued by “Autoridad Nacional de Licencias Ambientales ANLA”.

## Author contributions

MM and RIP conceived and, along with FJP and TLL, designed the study. CTGM, KGC, LAGG, SR, FJP, VP, MLV and TLL participated in field work and data collection. SR performed Symbiodiniaceae analysis. CTGM and TLL performed water optical properties analysis. TLL, CTGM, KGC and LAGG performed physiological analysis. TLL and RIP wrote the manuscript. MM, SR, CTGM, KGC and LAGG supported writing and editing the manuscript.

