## Supplementary Material for "Physiological and ecological consequences of the water optical properties degradation on reef corals"

López-Londoño et al.

#### Study site

Varadero Reef is located south-west of the Cartagena Bay in Colombia, close to the southern strait (Bocachica) that connects the Bay to the Caribbean Sea (Fig. 1). The reef has an extension of approximately 1 $km^2$ and appears to be a relic of a wider Plio-Pleistocene reef system (Díaz et al. 2000) that once dominated adjacent coastal areas (Lopez-Victoria et al. 2015; Pizarro et al. 2017). Coral cover is very low below 12 m. Above this depth, plate-like and small colonies typical of deep-environments become gradually abundant. Between 8 and 3 m coral cover is highest and massive colonies dominate the landscape. The shallow portion of the reef is still in good condition in terms of live coral cover (up to 50-60%) (Pizarro et al. 2017).

Rosario is part of a network of reticulated reefs occurring over the continental platform, being one of the largest coral reef areas in Colombia. In the vicinity of Rosario there is a large terrace with partially emerged portions and karstic depressions with exuberant coral growth. Massive colonies dominate in shallow waters and crustose and plate like morphologies gradually become dominant below 12 m. Coral cover has dramatically decreased during last decades throughout the reef system due to local, regional and global stressors (Díaz et al. 2000).

#### Chlorophyll a fluorescence: environmental conditions and PAM settings

Irradiances at the time of the effective quantum yield ($\Delta F/F_m'$) of photosystem II (PSII) measurements were 266.37 ± 78.61 and 100.69 ± 44.86 μmol quanta $m^{-2}$ $s^{-1}$ in Varadero and Rosario, respectively. $K_d$'s when measuring $\Delta F/F_m'$ in Varadero and Rosario were respectively 0.41 ± 0.08 $m^{-1}$ and 0.17 ± 0.01 $m^{-1}$.

Both for $\Delta F/F_m'$ and $F_v/F_m$ measurements taken respectively at noon and at dusk or dawn, a very weak beam of red light (<1 μmol quanta $m^{-2}$ $s^{-1}$) was used to estimate minimum fluorescence yields. Maximum fluorescence yields were determined after applying a short (800 ms) saturation pulse of actinic light (>4000 μmol quanta $m^{-2}$ $s^{-1}$). $\Delta F/F_m'$ and $F_v/F_m$ were calculated based on changes in fluorescence caused by the saturation pulse relative to the maximum fluorescence recorded at noon and dusk.

#### PE curves and photosynthetic parameters

To avoid potential artefacts related to different frequencies of dark / light periods on photosynthesis (Iluz 2012) when pulsating LEDs are used, light sources were controlled in continuous mode with a multifunction I/O card (USB-6001, National Instruments Corp., USA). The photosynthetic efficiency ($\alpha$), compensating irradiance ($E_c$), saturating irradiance ($E_k$), respiration rates ($R_d$), and maximum photosynthetic rates ($P_{max}$), were calculated from the light-limited and light-saturated regions of the PE curves following Iglesias-Prieto and Trench (1994) and Osinga (2012). The maximum quantum yield of oxygen evolution ($\Phi_{max}$) and the minimum quantum requirement ($1/\Phi_{max}$) were calculated based on the amount of light being absorbed and used to drive photosynthesis in the light-limited region of the PE curve. The ratio of absorbed light was calculated by multiplying the incident light spectrum and the *in-vivo* absorption spectra of corals calculated from reflectance measurements as $A = 1 - R$ (Schubert et al. 2011).

### Supplementary Figures

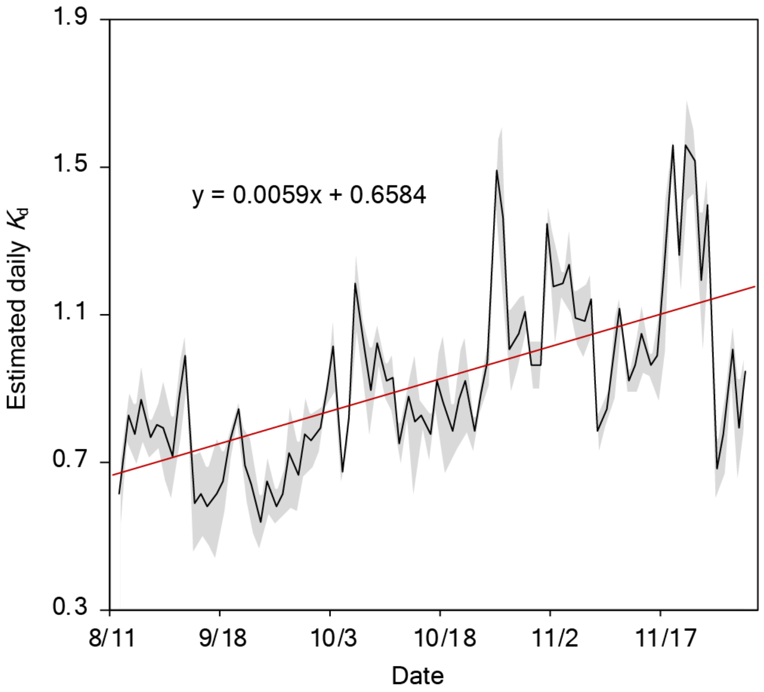


**Fig. S1.** Temporal variation of the mean daily $K_d$ in Varadero based on the synchronous oscillation of irradiances recorded by onshore and underwater light sensors (shaded area represents SD). The red line corresponds to the linear regression whose slope (m = 0.0059) was used for correcting the cumulative signal attenuation of the sensors.

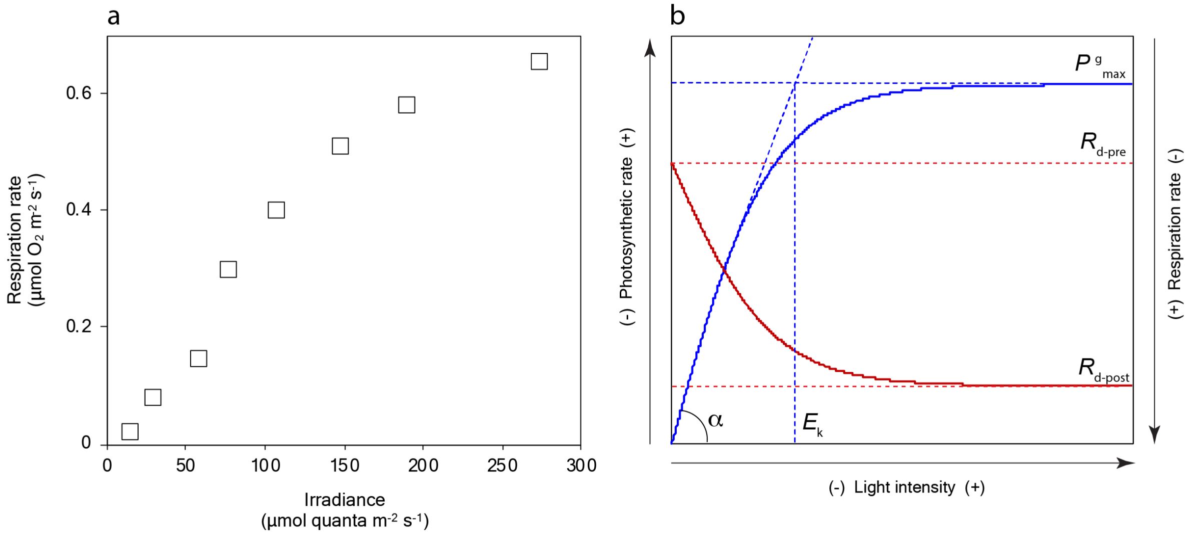


**Fig. S2.** Photosynthesis and respiration of the coral holobiont as a function of irradiance. **a** Respiration rates of *O. faveolata* recorded immediately after exposing corals to a series of increasing light levels. Measurements were performed under laboratory conditions using corals from the Puerto Morelos lagoon, Mexico, held in outdoor aquariums at ~28 °C and 50% incident irradiance. This data, together with evidence from other studies (Colombo-Pallotta et al. 2010; Holcomb et al. 2014), indicate that there is a light-associated asymptotic increase of the respiratory activity until reaching a maximum potentially determined by the light-saturated photosynthetic rates ($E_k$). **b** Schematic representation of the variation of the photosynthetic (blue line) and respiration (red line) rates as a function of irradiance. These parameters, which were calculated with hyperbolic tangent functions, were used to quantify the daily phototropic contribution of algal symbionts to the energy requirements of *O. faveolata*. $P^g_{max}$ is the maximum gross photosynthetic rate, $\alpha$ is the photosynthetic efficiency, $R_{d\text{-}pre}$ and $R_{d\text{-}post}$ are respectively the pre- and post-illumination respiration rates, and $E_k$ is the compensating irradiance.

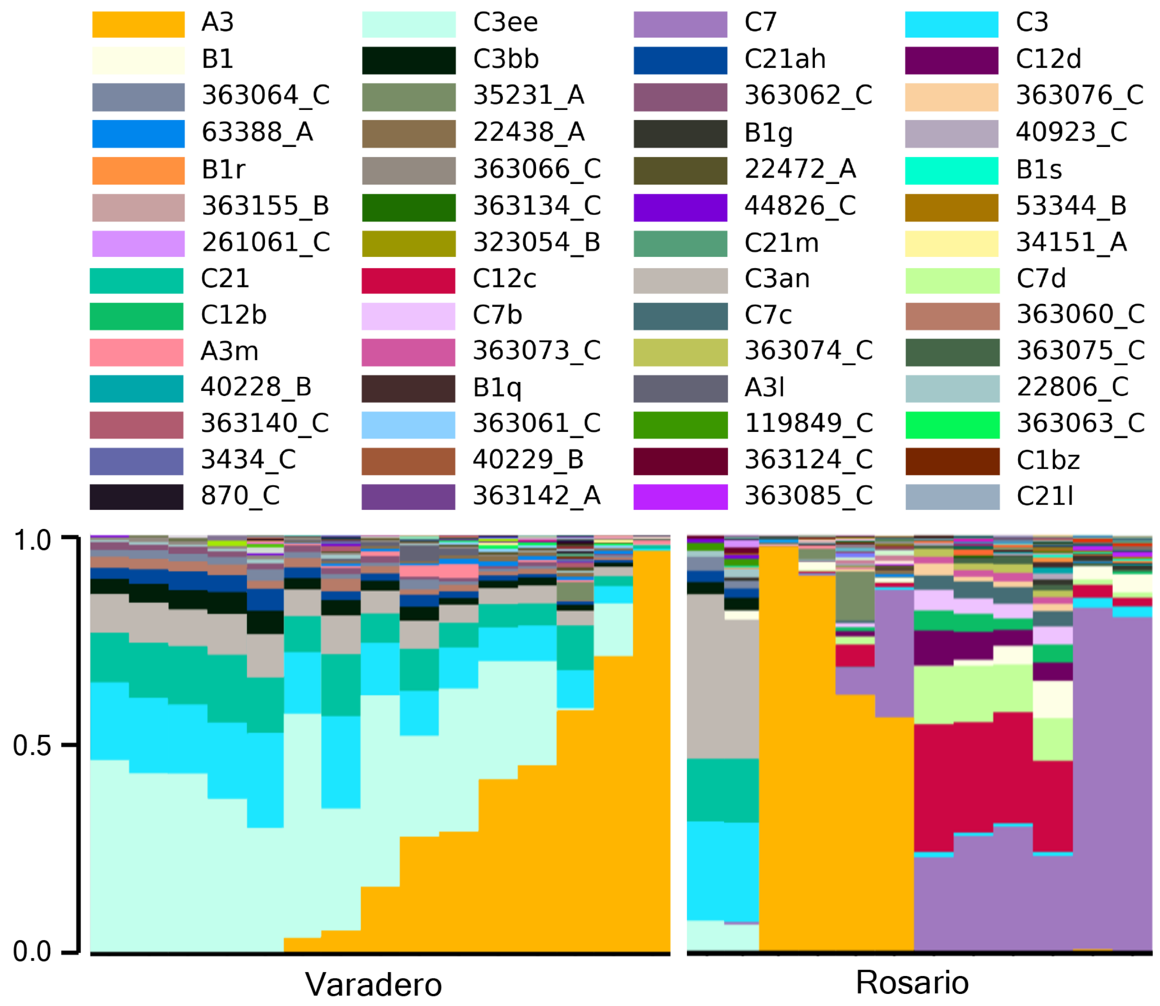


**Fig. S3.** Raw diversity of Symbiodiniaceae ITS2 type profiles in coral samples from Varadero and Rosario. Each column of stacked bars represents one sample, with the bar heights within each column representing the relative abundance of a given sequence in the sample.

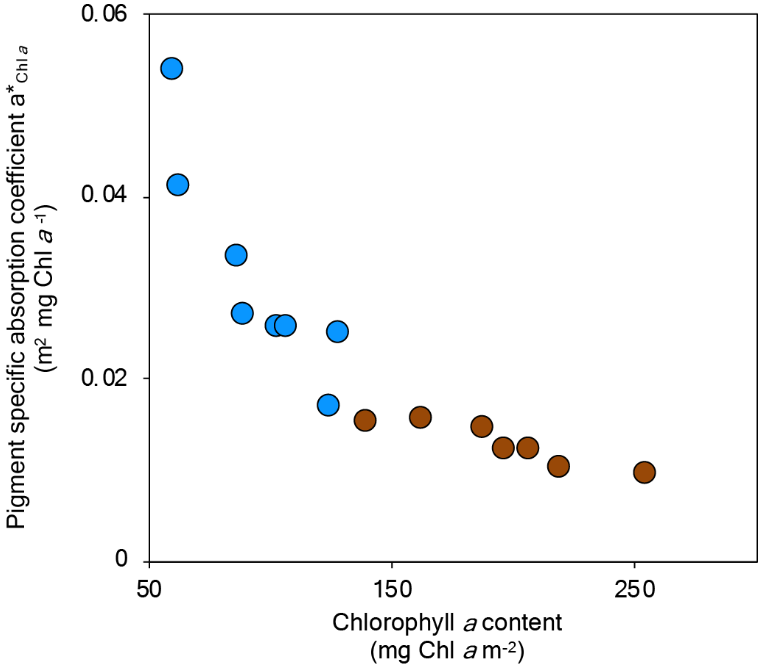


**Fig. S4.** Changes in the pigment specific absorption coefficient ($a^*_{Chl\ a}$) as a function of the chlorophyll *a* content in *Orbicella faveolata* corals from Varadero at 3.5 m (brown circles) and Rosario at 12 m (blue circles).

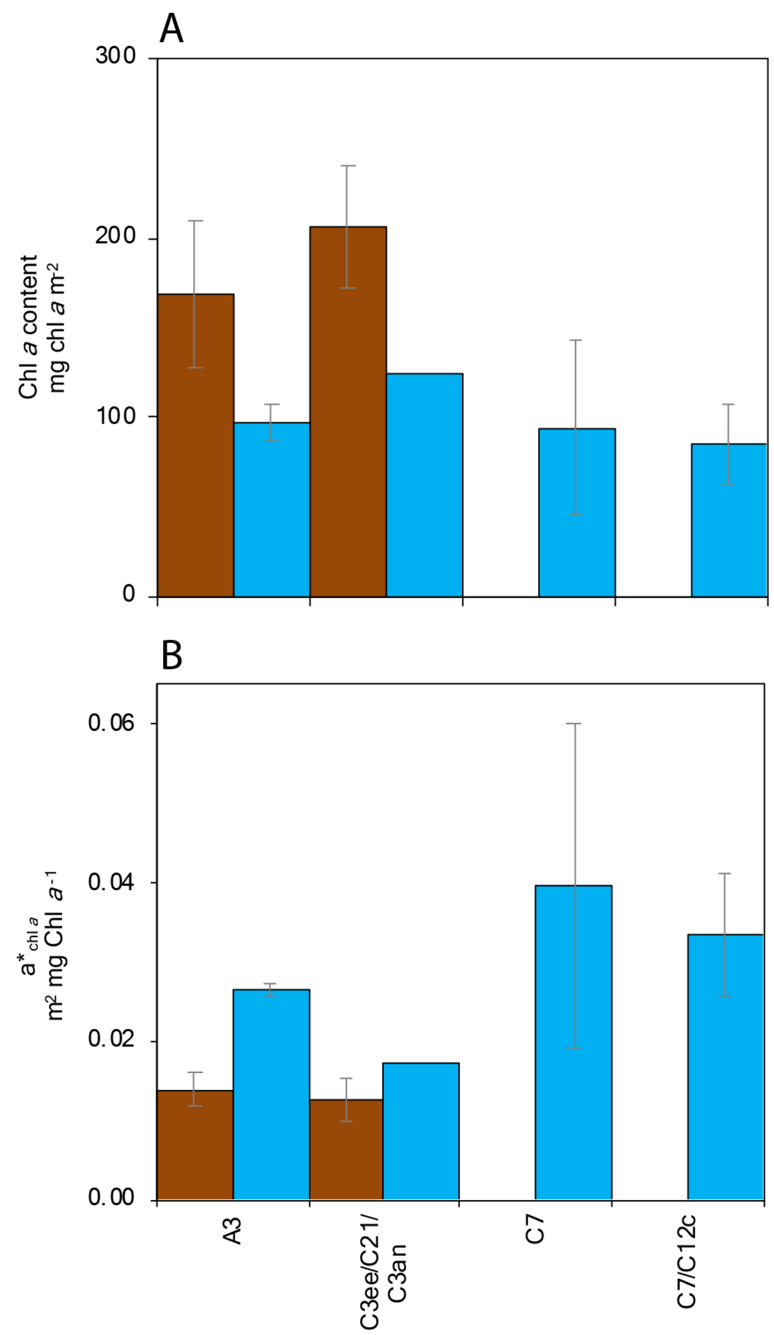


**Fig. S5.** Variation in chlorophyll *a* content and the pigment specific absorption coefficient ($a^*_{Chl\ a}$) as a function of the dominant symbionts in coral samples from Varadero (brown columns) and Rosario (blue columns).

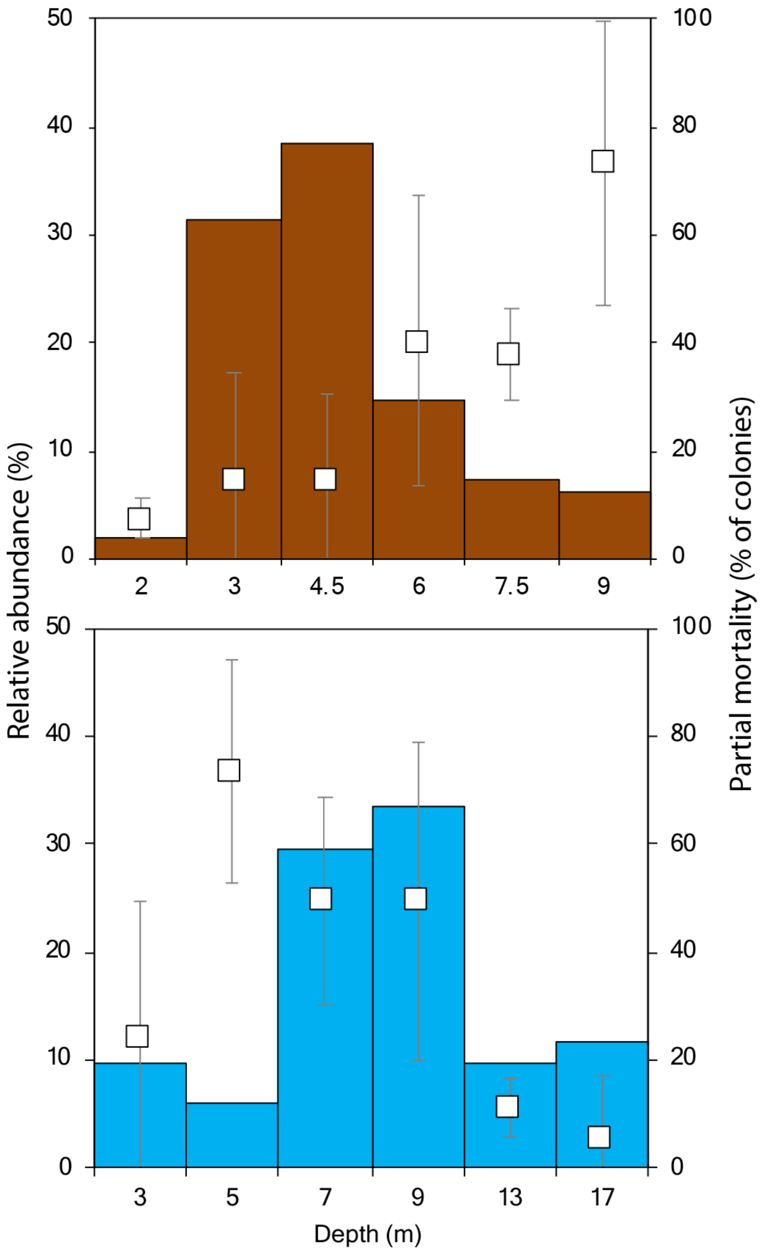


**Fig. S6.** Variation in relative abundance and percentage of old mortality of *O. faveolata* colonies across depths. Top panel: Varadero reef; lower panel: Rosario reef. Values of old mortality correspond to the mean ± SD.

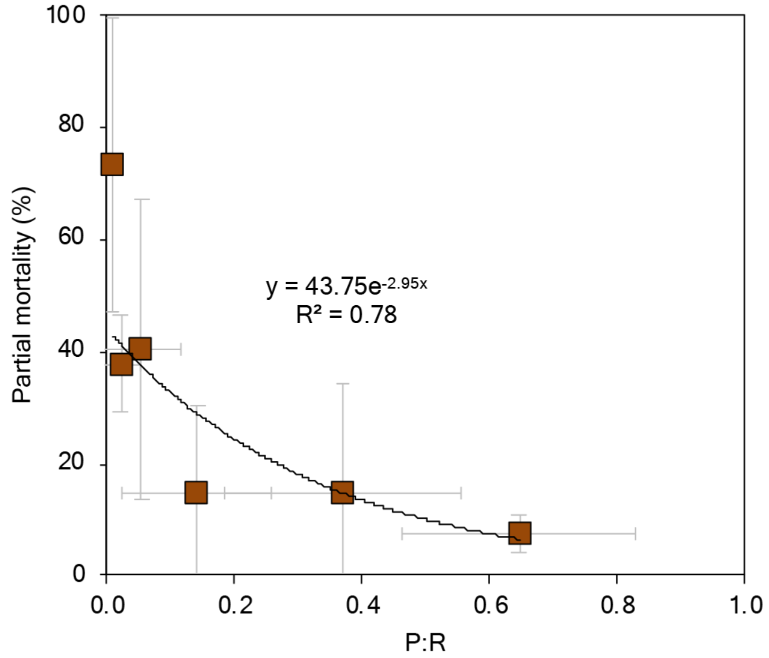


**Fig. S7.** Partial mortality of *O. faveolata* colonies as a function of the integrated daily P:R ratios in Varadero. Values correspond to the mean ± SD. An exponential regression was used to fit the data (y = 43.75e-2.95x).
